# Globally defining the effects of mutations in a picornavirus capsid

**DOI:** 10.1101/2020.10.06.327916

**Authors:** Florian Mattenberger, Victor Latorre, Omer Tirosh, Adi Stern, Ron Geller

## Abstract

The capsids of non-enveloped viruses are highly multimeric and multifunctional protein assemblies that protect the viral genome between infection cycles, dictate host and cell tropism, and mediate evasion of humoral immune responses. As such, capsids play key roles in viral biology and pathogenesis. Despite their importance, a comprehensive understanding of how mutations affect viral fitness across different structural and functional attributes of the capsid is lacking. To address this limitation, we globally define the effects of mutations in the capsid of a human picornavirus, generating a comprehensive dataset encompassing >90% of all possible single amino acid mutations. Moreover, we use this information to identify structural and sequence determinants that accurately predict mutational fitness effects, refine evolutionary analyses, and define the sequence specificity of key capsid encoded motifs. Finally, capitalizing on the sequence requirements identified in our dataset for capsid encoded protease cleavage sites, we implement and validate a bioinformatic approach for identifying novel host proteins targeted by viral proteases. Our findings present the most comprehensive investigation of mutational fitness effects in a picornavirus capsid to date and illuminate important aspects of viral biology, evolution, and host interactions.

## INTRODUCTION

The capsids of non-enveloped viruses are among the most complex of any viral protein. These highly multimeric structures must correctly assemble around the genome from numerous subunits, at times numbering in the hundreds, while avoiding aggregation ^1–3^. Moreover, the assembled structure must be both sufficiently stable to protect the viral genome during its transition between cells yet readily disassemble upon entry to initiate subsequent infections. For these functions to be achieved, viral capsids must encode the information for interacting with numerous cellular factors that are required to correctly fold and assemble around the genome ^4–8^. Viral capsids also play key roles in pathogenesis, dictating host and cell tropism by encoding the determinants for binding cellular receptors ^9,10^ and mediating escape from humoral immune responses ^11,12^. As a result, viral capsids show the highest evolutionary rates among viral proteins.

The picornaviruses constitute a large group of single-stranded, positive-sense RNA viruses and include several pathogens of significant medical and economic impact ^13^. Their relative simplicity and ease of culture have made picornaviruses important models for understanding virus biology. Among the many breakthroughs achieved with these viruses was the determination of the first high-resolution structure of the capsid of an animal virus, making the picornavirus capsid the prototypical non-enveloped, icosahedral viral capsid ^13^. Picornavirus capsid genesis initiates with the co-translational release of the P1 capsid precursor protein from the viral polyprotein via the proteolytic activity of the viral encoded 2A protease ^7,13^. Subsequently, the viral encoded 3CD protease (3CD^pro^) cleaves the P1 capsid precursor to liberate three capsid proteins (VP0, VP3, and VP1), generating the capsid protomer. Five protomers then assemble to form the pentamer, twelve of which assemble around the viral genome to yield the virion. Finally, in some picornaviruses, VP0 is further cleaved into two subunits, VP4 and VP2, following genomic encapsidation to generate the infectious, 240 subunit particle ^7,13^. Work over the years has identified numerous host factors that help support capsid formation ^4,5,14–16^, defined antibody neutralization sites ^11^, and identified numerous host receptors for many members of this viral family ^9^.

Despite significant progress in understanding the structure and function of picornavirus capsids, a comprehensive understanding of how mutations affect viral fitness across different structural and functional attributes is lacking. To address this, we perform a comprehensive analysis of mutational fitness effects (MFE) across the complete capsid region of the human picornavirus coxsackievirus B3 (CVB3), analyzing >90% of all possible single amino acid mutations. Furthermore, using this data, we develop models to predict the effect of mutations with high accuracy from available sequence and structural information, improve evolutionary analyses of CVB3, and define the sequence preferences of several viral encoded motifs. Finally, we use the information obtained in our dataset for the sequence requirements of capsid encoded 3CD protease cleavage sites to identify host targets of this viral protease. Overall, our data comprise the most comprehensive survey of MFE effects in a picornavirus capsid to date and provide important insights into virus biology, evolution, and interaction with the host.

## RESULTS

### Deep mutational scanning of a CVB3 capsid

To generate CVB3 libraries encoding a large amount of diversity in the capsid region, we used a codon-level PCR mutagenesis method ^17^. The mutagenesis protocol was performed on the capsid precursor region P1 in triplicate to generate three independent mutagenized libraries (Mut Library 1-3; Fig. 1A). From these, three independent viral populations (Mut Virus 1-3) were derived by electroporation of in vitro transcribed viral RNA into HeLa-H1 cells (Fig. 1A). High-fidelity next-generation sequencing ^18^ was then used to analyze the mutagenized libraries and resulting viruses, unmutagenized virus populations (WT virus 1-2), as well as controls for errors occurring during PCR (PCR) and reverse transcription (RT-PCR). High coverage was obtained for all samples (>10^6^ per codon across all experimental conditions and >6.5×10^5^ for the controls; Supplementary Table S2). Due to the high rate of single mutations within codons observed in the RT-PCR control compared to the mutagenized virus populations (Supplementary Table S2), all single mutants were omitted from our analysis to increase the signal-to-noise ratio. While this resulted in an inability to analyze 83.4% of synonymous codons in the capsid region (1746/2094) only 2.8% of non-synonymous mutations were lost to analysis (458/16,169). Upon removing single mutations within codons, we obtained a large signal-to-noise ratio in the average mutation rate of 510x (range 449–572) and 245x (range 174–285) for the mutagenized libraries and viruses, respectively, compared to their error controls (Fig. 1B and Supplementary Table S2). On average, 0.9 (range 0.8–1.02) codon mutations were observed per genome, which was in agreement with Sanger sequencing of 59 clones (range 18–23 per library; Fig. S1 and Supplementary Table S3). As expected, the rate of stop codons, which should be invariably lethal in the CVB3 capsid, decreased significantly following growth in cells to <0.5% of that observed in the corresponding mutagenized libraries (p < 0.005 by paired t-test on log-transformed data; Supplementary Table S2). No major bias was observed in the position within a codon where mutations were observed (Fig. S2A) nor in the type of mutation (Fig. S2B), except for the WT virus, which had a high rate of A to G transitions in the two independent replicates analyzed. Of all 16,169 possible amino acid mutations in the capsid region (851 AA x 19 AA mutation = 16,169), a total of 14,839 amino acid mutations were commonly observed in all three mutagenized libraries, representing a 91.8% of all possible amino acid mutations in the capsid region, allowing us to globally assess the effects of the vast majority of amino acid mutations on the capsid (Fig. 1C).

**Figure 1.**
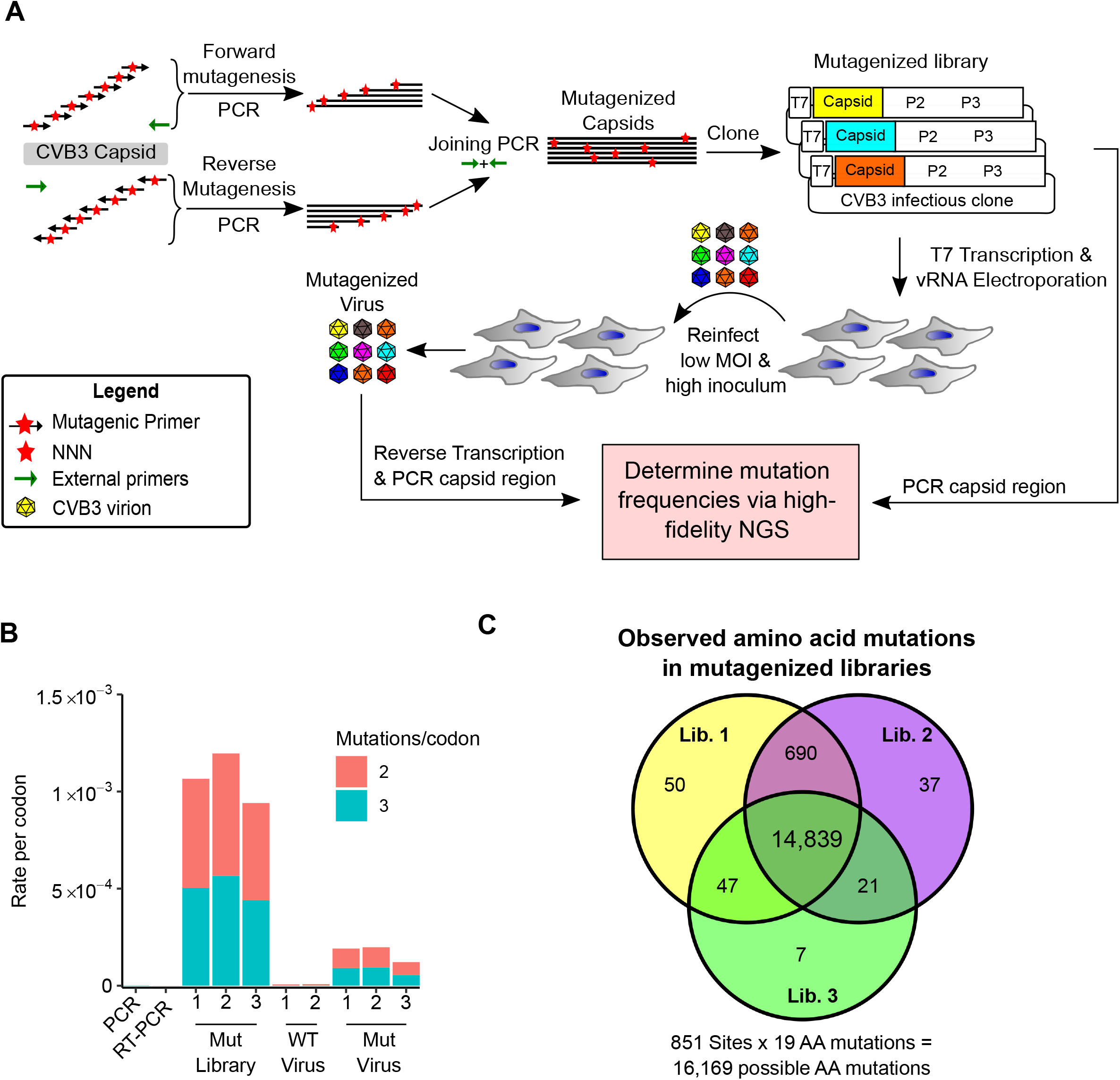
Deep mutational scanning (DMS) of the CVB3 capsid. **A.** Overview of the deep mutational scanning experimental approach. A mutagenesis PCR was performed using an external reverse primer and a pool of forward mutagenic primers targeting each capsid codon that encode degenerate nucleotides (NNN) at the codon matching position. Similarly, a reverse mutagenesis PCR reaction was performed. The products of these PCRs were joined using the external primers and cloned into a CVB3 infectious clone to generate the mutagenized libraries. This process was performed in triplicate, generating 3 libraries (Mut Library 1-3). Viral genomic RNA (vRNA) produced from the mutant libraries was electroporated into cells to generate high diversity CVB3 populations (Mut Virus 1-3). The relative frequency of each mutation relative to the WT amino acid was then determined in both the mutagenized libraries and the resulting virus populations via high-fidelity duplex sequencing. **B.** The average rate of double or triple mutations per codon observed in the mutagenized libraries (Mut Library 1-3), the resulting mutagenized virus (Mut Virus 1-3), as well as controls for the error rate of the amplification and sequencing process (PCR and RT-PCR) or the WT unmutagenized virus (WT Virus 1-2). Single mutations per codon were omitted from the analysis to increase the signal-to-noise ratio. **C.** Venn diagram showing the number of amino acid mutations observed in the mutagenized libraries.

### Mutational fitness effects across the CVB3 capsid

We next derived the mutational fitness effects (MFE) of each observed mutation by examining how its frequency changed relative to that of the WT sequence following growth in cells. The preferences for the different amino acids at each position (amino acid preferences ^19^) showed a high correlation between biological replicates (Spearman’s ρ > 0.83; Supplementary Figure S3 and Supplementary Table S4 MFE). Overall, most mutations in the capsid were deleterious, with only 1.2% of mutations increasing fitness relative to the WT amino acid (Fig. 2A and Supplementary Table S4). Hotspots where mutations were tolerated were observed at several regions across the capsid (Fig. 2A). These hotspots largely overlapped with highly variable regions in natural sequences, as measured by Shannon entropy in the enterovirus B family, indicating that lab measured MFE reflect natural evolutionary processes (Fig. 2A, top). Indeed, a strong correlation was observed between MFE and sequence variability for the enterovirus B genus (Spearman’s ρ = 0.59, p < 10^−16^; Fig. 2B). Similarly, antibody neutralization sites overlapped with hotspots for mutations (Fig. 2A, top) and were significantly less sensitive to mutations (p < 10^−16^ by Mann-Whitney test; Fig. 2C). As expected, mutations were also less deleterious in loops compared to β-strands (p < 10^−16^ by Kruskal-Wallis test; Fig. 2D), at surface residues compared to core residues (p < 10^−16^ by Kruskal-Wallis test; Fig. 2E), and for mutations predicted to be destabilizing (p < 10^−16^ by Mann-Whitney test; Fig. 2F) or aggregation-prone (p < 10^−16^ by Mann-Whitney test; Fig. 2G). Importantly, independent validation of the MFE of 10 different mutants using a sensitive qPCR method ^20^ showed a strong correlation with the DMS results (Spearman’s ρ = 0.9, p < 0.001; Supplementary Table S5).

**Figure 2.**
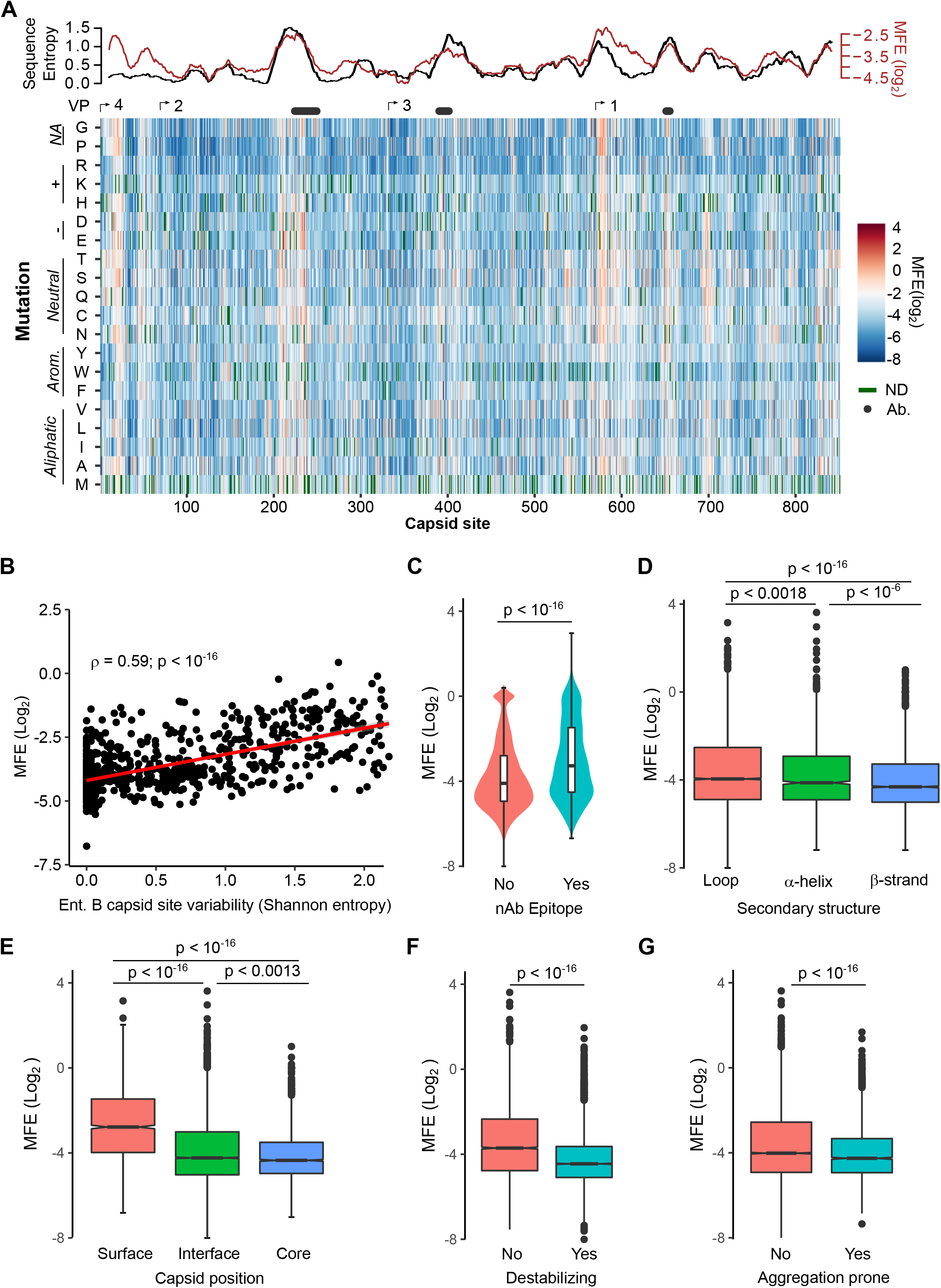
Mutational fitness effects across the CVB3 capsid and their correlation with structural, evolutionary, and immunological attributes. **A.** Overview of the mutational fitness effects (MFE) across the CVB3 capsid. Bottom: a heatmap representing the MFE of all mutations observed at each capsid site. Green indicates no data available (ND), and the positions of the mature viral proteins (VP1-4) or antibody neutralization sites (nAb) are indicated above. Top: A 21 amino acid sliding window analysis of the average sequence variation in CVB3 genomes (Shannon entropy; black line) or the average MFE (red line). **B.** Correlation between derived MFE and variation in enterovirus B sequence alignments (Shannon entropy). **C.** Violin plot of MFE in antibody neutralization sites versus other capsid sites. **D-G**. Boxplots of MFE as a function of secondary structure (D), position in the capsid (E), or the predicted effect of mutations on stability (F) or aggregation propensity (G). Two-sided Mann-Whitney or Kruskal-Wallis tests were used for 2 or 3 category comparisons, respectively.

### Prediction of MFE from available structural and sequence information

As MFE correlated with natural sequence variation and different structural features of the capsid (Fig. 2), we next investigated if MFE could be predicted from available structural and sequence information. For this, we obtained a dataset of 52 parameters, including structural information derived from the crystal structure of the CVB3 capsid (PDB:4GB3), amino acid properties, natural variation in available enterovirus sequences (Shannon entropy), and predicted the effects of mutation on stability and aggregation propensity using FoldX ^21^ and TANGO ^22^, respectively (Supplementary Table S6). We then employed a random forest algorithm to identify the parameters that can best predict MFE, limiting our analysis to sites that present in the crystal structure and where mutations were observed in at least 2 replicates to improve accuracy (total of 9,685 mutations). Overall, a model trained on 70% of the dataset was able to predict the remaining 30% of the data (2,905 mutations) with high accuracy (Spearman’s ρ > 0.75, Pearson’s r = 0.76; p < 10^−16^; Fig. S4A,B). Surprisingly, a random forest model trained on the top five predictors alone showed similar accuracy (Spearman’s ρ = 0.73, Pearson’s r = 0.73; p < 10^−16^; Fig. 3B). Excluding natural sequence variation, amino acid identity, or structural attributes reduced model predictability significantly (>20%; data not shown), suggesting a combination of evolutionary, sequence, and structural information best explains MFE. Using an alternative approach, we were able to predict the data with slightly lower accuracy using a linear model with the same five predictors (p < 10^−16^, Spearman’s ρ = 0.67, Pearson’s r = 0.67; Fig. S4C). Together, these results suggest that the prediction of MFE in the CVB3 capsid can be achieved at relatively high accuracy based on available structural and sequence information. Due to the high conservation of capsid structure in picornaviruses, as well as the availability of numerous capsid sequences and structures, these findings are likely generalizable to related picornaviruses.

**Figure 3.**
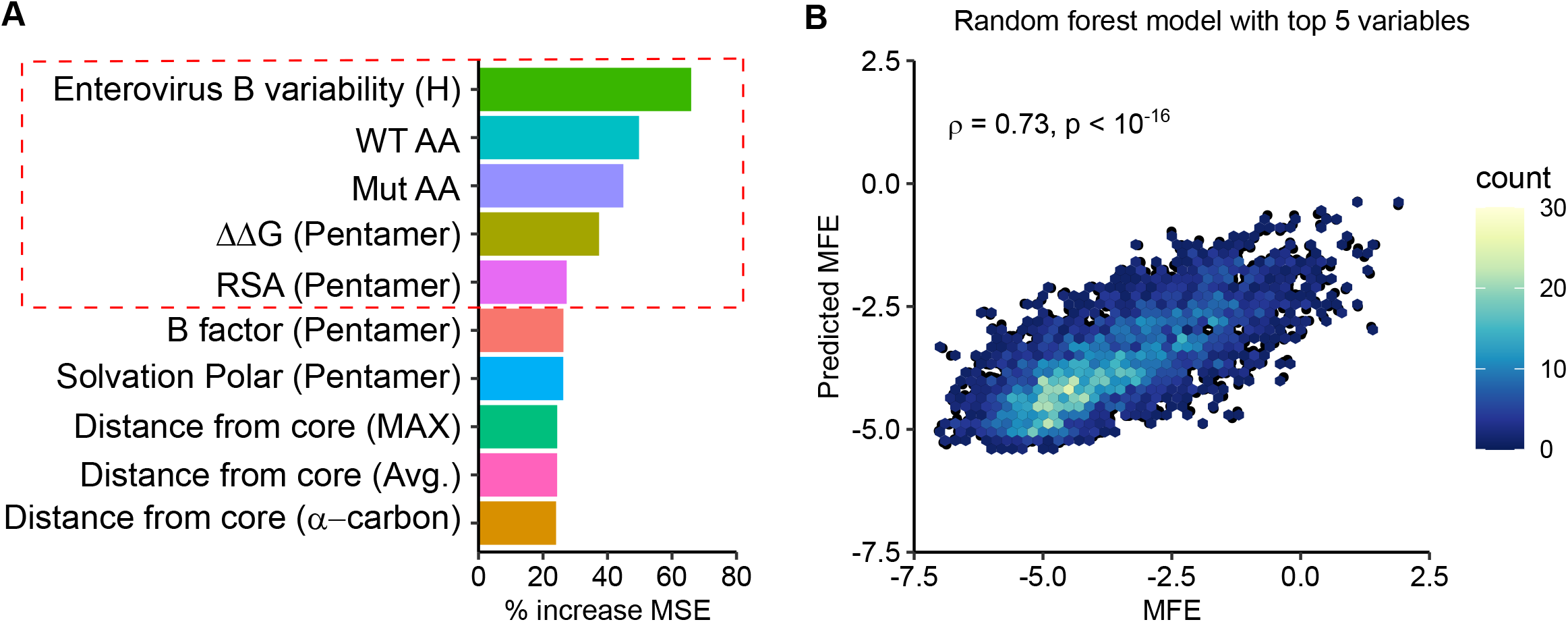
Prediction of MFE based on structural and sequence information. **A.** The top 10 predictors identified in a random forest model for explaining MFE in the CVB3 capsid based on the percent of mean squared error (MSE) increase. **B.** Hexagonal plot showing the correlation between MFE predicted using a random forest algorithm trained on the top 5 variables versus observed MFE. The random forest model was trained on 70% of the data, and then tested on the remaining 30% (shown). RSA, relative surface area.

### Experimentally measured MFE inform of natural evolutionary processes

We next examined if our experimentally measured MFE could improve phylogenetic models of CVB3 evolution by incorporating site-specific amino acid preferences using PhyDMS ^23^. Indeed, significant improvement in model fit was observed (Table 1 PHY; p < 10^−16^ using a log-likelihood test compared to non-site-specific codon models), supporting the relevance of our results to understanding evolutionary processes in nature. Nevertheless, selection in nature was significantly more stringent than in the lab (β= 2.18), indicating the presence of additional selection pressures. As laboratory conditions lack selection from antibodies, we used the sum of the absolute differential selection observed at each site ^24^ to examine whether known antibody neutralization sites show differential selection between the two environments (Supplementary Table S7). Indeed, antibody neutralization sites showed significantly higher differential selection values compared to other residues (p < 10^−6^ by Mann-Whitney test; Fig. 4A). Moreover, the three sites showing the strongest overall differential selection were found in known antibody neutralization sites: position 226 and 242 in the EF loop (residues 157 and 173 of VP2) and position 650 in the BC loop (residue 80 of VP1; Fig. 4B-D and Supplementary Table S7). In summary, incorporation of our experimentally derived amino acid preferences into phylogenetic analyses significantly improved model fit and identified residues in antibody neutralization sites that show differential selection, suggesting these may play important roles in immune evasion in vivo.

**Table 1:** Incorporation of DMS results in evolutionary models better describes natural CVB3 evolution compared to standard codon models.

| Model | $\Delta AIC$ | LogLikelihood | Parameters | Parameter Values |
| --- | --- | --- | --- | --- |
| ExpCM | 0.00 | -14580.51 | 6 | beta=2.18, kappa=7.47, omega=0.16 |
| Goldman-Yang M5 | 4187.56 | -16668.29 | 12 | alpha_omega=0.30, beta_omega=10.00, kappa=7.15 |
| Averaged ExpCM | 4303.74 | -16732.38 | 6 | beta=0.61, kappa=7.55, omega=0.02 |
| Goldman-Yang M0 | 4371.26 | -16761.14 | 11 | kappa=7.14, omega=0.02 |

**Figure 4.**
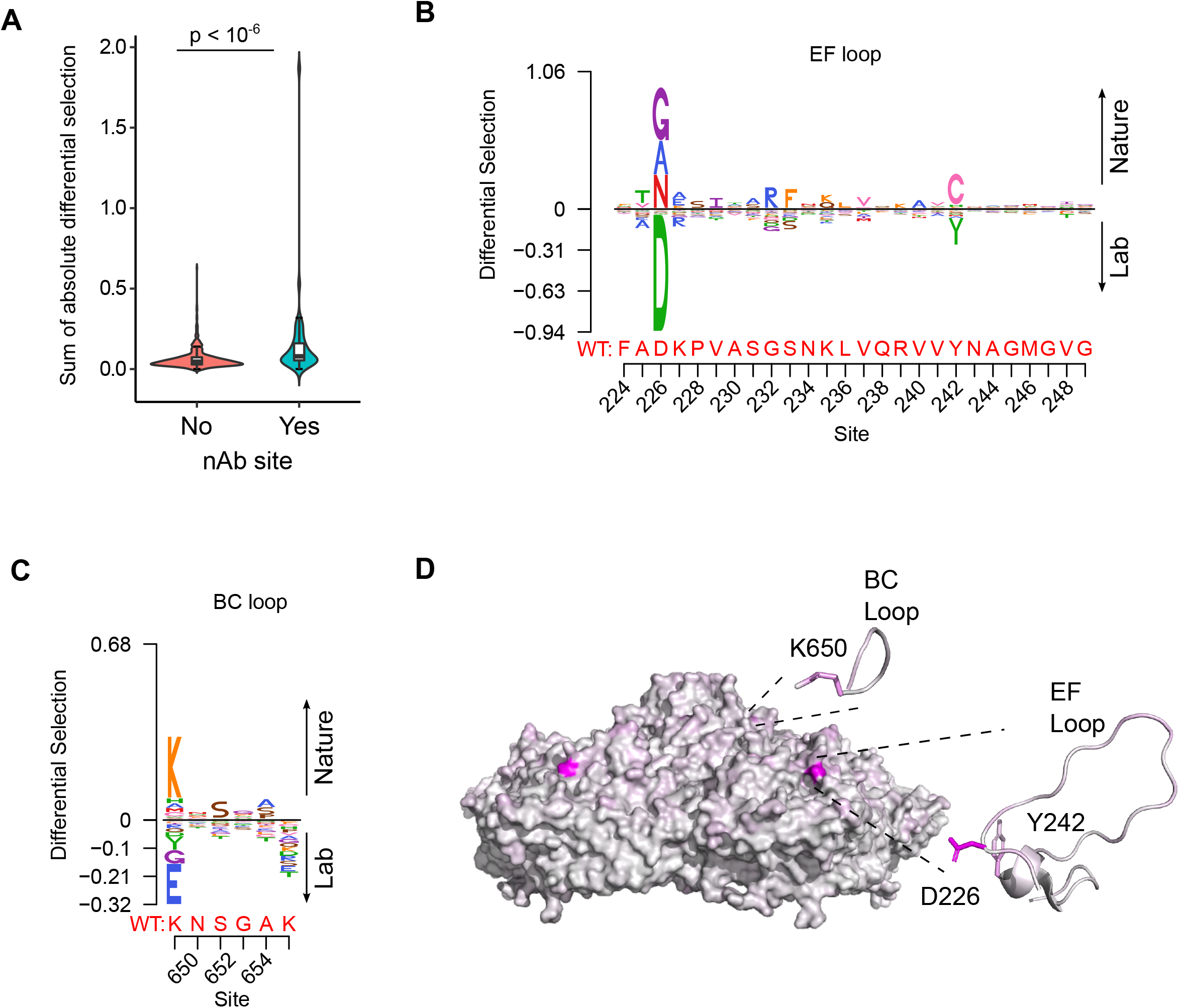
Antibody neutralization sites show differential selection between laboratory conditions and nature. **A.** Violin plot showing the sum of absolute differential selection observed at capsid sites comprising antibody neutralization epitopes (nAb) versus all other capsid sites. **B-C.** Logoplots showing the observed differential selection of sites in the EF loop or BC loop. The WT sequence is indicated in red. **D.** The CVB3 capsid pentamer (PDB:4GB3), colored according to the amount of differential selection. The BC and EF loops are shown next to the structure together with the sidechains for sites showing the highest differential selection.

### Insights into capsid encoded motifs: Myristoylation and protease cleavage

Picornavirus capsids undergo a complex assembly path to generate the infectious particle. These include myristoylation, cleavage by the viral proteases 2A and 3CD^pro^, as well as interaction with cellular chaperones and glutathione ^4,7,14–16^ (Fig. 5A). Having obtained a comprehensive dataset for MFE across the capsid, we next examined the sequence requirements for several of these capsid encoded motifs. Specifically, myristoylation of the N-terminal glycine is essential for virion assembly ^16^. In agreement with this, the N-terminal glycine in the CVB3 capsid showed the strongest average fitness cost upon mutation in the capsid (Fig. S5 and Supplementary Table S4). The remaining sites in the myristoylation motif agreed with the canonical myristoylation motif in cellular proteins (Prosite pattern PDOC00008) ^25^, albeit with increased selectivity at three of the six positions (Fig. S5A). On the other hand, a conserved WCPRP motif in the C-terminal region of VP1 that was shown to be important for 3CD^pro^ cleavage of the related foot and mouth disease virus capsid (FDMV; YCPRP motif) ^26^ was found to be intolerant to mutations compared to other capsid residues (p < 0.05 versus all other positions by Mann-Whitney test; sites 815-819 in CVB3). Moreover, within this motif, the sites showing the highest average fitness cost in our DMS dataset were identical to analogous positions in FMDV that resulted in a loss of viability upon mutation to alanine (Fig. S5B) ^26^, highlighting the conservation of this motif across different picornaviruses.

**Figure 5.**
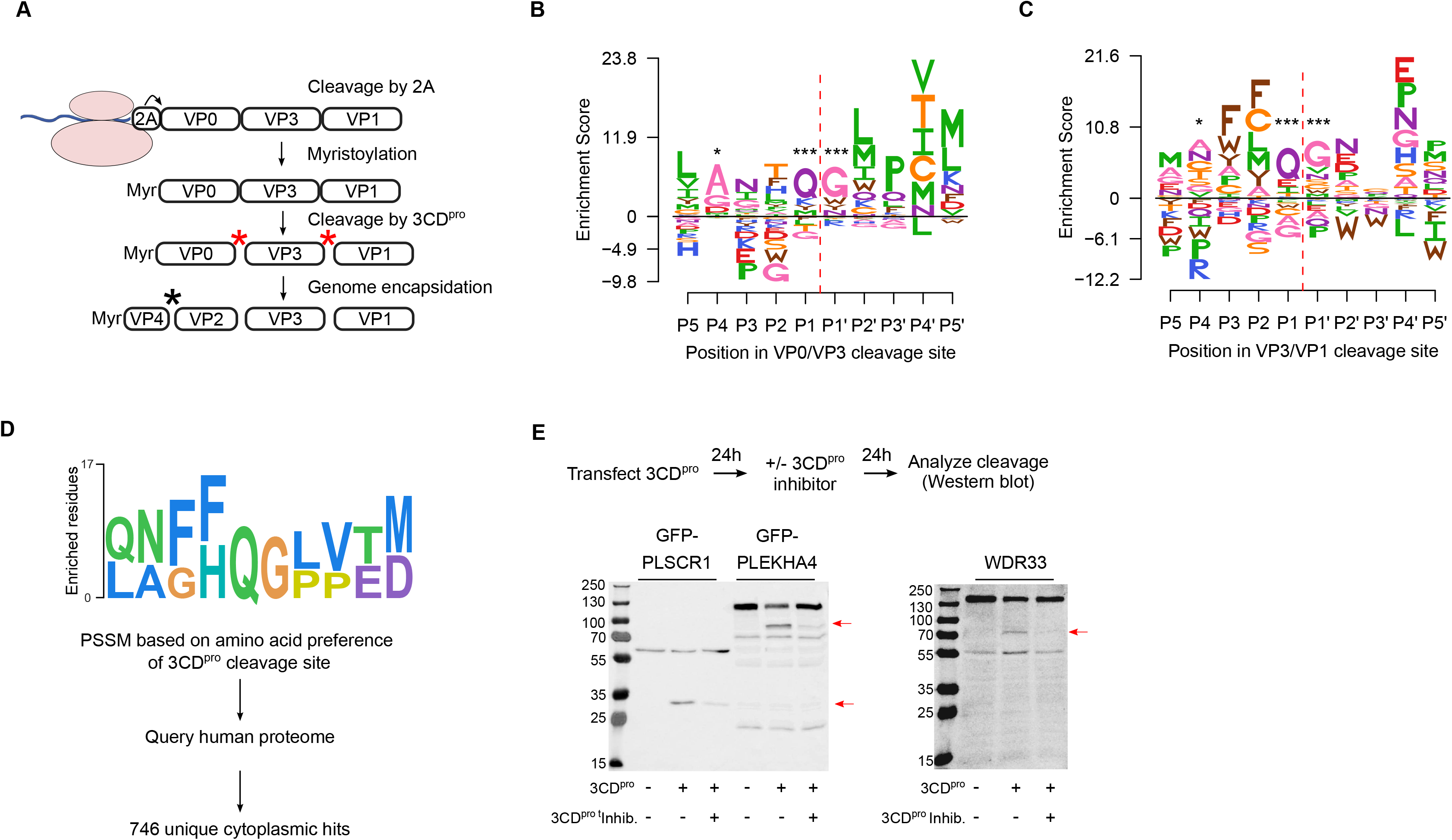
Sequence preference of capsid 3CD^pro^ cleavage sites and their use for the identification of novel cellular targets of the viral protease. **A.** Overview of the CVB3 capsid maturation pathway. The CVB3 capsid precursor P1 is co-translationally cleaved by the viral 2A protease. P1 is then myristoylated and cleaved by the viral 3CD^pro^ to generate the capsid proteins VP0, VP3, and VP1. Finally, upon assembly and genome encapsidation, VP0 is further cleaved into VP4 and VP2 in a protease independent manner to generate the mature capsid. Red and black astrickses indicated 3CD^pro^ or protease independent cleavage events, respectively. **B-C.** Logoplots showing amino acid preferences for the 10 amino acid region spanning the 3CD^pro^ cleavage sites (P1-P’1) of both VP0/VP3 and VP3/VP1 in the DMS dataset. **D.** Overview of the bioinformatic pipeline for identification of novel 3CD^pro^ cellular targets using the amino acid preferences for the capsid cleavage sites from our DMS study. A position-specific scoring matrix (PSSM) was generated based on the amino acid preferences for the 10 amino acid region spanning the two 3CD^pro^ cleavages sites. This PSSM was then used to query the human genome for potential cellular targets, and non-cytoplasmic proteins were filtered out, yielding 746 proteins. **E.** The cellular proteins PLSCR1, PLEKHA4, and WDR33 are cleaved by 3CD^pro^. Western blot analysis of cells cotransfected with 3CD^pro^ and GFP-PLSCR1 or GFP-PLEKHA4 and probed with a GFP antibody or transfected with 3CD^pro^ and probed using a WDR33 antibody. When indicated, the 3CD^pro^ inhibitor rupintrivir was included to ensure cleavage was mediated by the viral protease. Red arrows indicate cleavage products of the expected size (GFP-PLSCR1 full length = 64 kDa, cleaved N-terminus = 36 kDa; GFP-PLEKHA4 full length = 118, cleaved N-terminus = 72 kDa; WDR33 full length = 146 kDa, cleaved N-terminus = 72 kDa). * p < 0.05, *** p< 0.001.

The viral 3C protease (3C^pro^) cleaves the picornavirus capsid at two conserved glutamine-glycine (QG) pairs to liberate the viral capsid proteins VP0, VP3, and VP1 (Fig. 5A). Previous work has defined the sequence specificity of several picornavirus 3C^pro^ enzymes by examining both natural sequence variation and in vitro cleavage assays using synthetic peptides ^27^. However, unlike other 3C^pro^ mediated cleavage events in the viral polyprotein, the capsid is only efficiently cleaved by the precursor protein 3CD^pro^ ^28^. To gain insights into the sequence specificity of 3CD^pro^, we examined the amino acid preferences for a 10 amino acid region surrounding the protease cleavage site (P5-P5’). As expected based on the known specificity of the 3C protease ^27^, a strong preference for the presence of QG was observed at both 3CD^pro^ cleavage sites in our dataset (positions P1 and P1’ in the cleavage site; Fig. 5B,C). Interestingly, significant correlation in amino acid preferences between the two cleavage sites was observed only at P1-P1’ (Pearson’s ρ > 0.99, p < 10^−16^) and P4 (Pearson’s ρ > 0.49, p < 0.05), as was the case in the enterovirus B alignments (Pearson’s ρ > 0.84 and p < 10^−6^ for positions P4, P1, and P’1; data not shown). Hence, the low agreement in amino acid preferences observed for most positions across the two 3CD^pro^ cleavage sites suggests cleavage is strongly dictated by positions P4, P1, and P1’.

### Identification of 3CD^pro^ cellular targets based on the sequence preferences of capsid encoded protease cleavage sites

In addition to cleaving the viral polyprotein, the picornavirus proteases cleave cellular factors to facilitate viral replication, including both antiviral factors and cellular factors that favor viral IRES-driven translation mechanism over cellular cap-dependent translation (e.g. DDX58, eIF4G, and PABP) ^27,29^. As the canonical 3C/3CD^pro^ QG cleavage site occurs on average 1.6 times per protein in the human proteome (~33,000 times), we sought to examine if the rich dataset we obtained for the amino acid preferences of the capsid 3CD^pro^ cleavage sites can be used to identify novel cellular factors that are targeted by the viral protease. Specifically, a position-specific score matrix (PSSM) was generated for the 10 amino acid region spanning the two protease cleavage sites in the CVB3 capsid (P5-P5’) based on the amino acid preferences identified in our study (Fig. 5D). This PSSM was then used to query the human proteome for potential cleavage sites, yielding a total of 746 cytoplasmic proteins (Fig. 5D; Supplementary Table S8). Eleven cellular factors that are known to be cleaved during enterovirus infection were identified using this approach, including the viral sensor Probable ATP-dependent RNA helicase DDX58 (RIG1), the immune transcription factors p65 (RELA) and interferon regulatory factor 7 (IRF7), and polyadenylate-binding protein 1 (PABPC1), an important factor in translation initiation and mRNA stability (Supplementary Table S8) ^27,30^.

To evaluate whether our approach can identify novel cellular targets for the viral protease, we examined the ability of 3CD^pro^ to cleave eight different proteins found in the data set, focusing on those with cellular functions of potential relevance to CVB3 biology and which could be readily detected in our cell culture assay (e.g. availability of antibodies or tagged-variants, cleavage fragments of observable size, and high expression level). These included four interferon-inducible proteins (Pleckstrin Homology Domain Containing A4, PLEKHA4; Phospholipid Scramblase 1, PLSCR1; NOD-like receptor family CARD domain containing 5, NLRC5; Zinc Finger CCCH-Type Containing, Antiviral 1, ZC3HAV1) and four proteins involved in various cellular functions, namely apoptosis (MAGE Family Member D1, MAGED1), RNA processing (WD repeat domain 33, WDR33), and vesicle transport (Cyclin G Associated Kinase, GAK; Tumor Susceptibility 101, TSG101). Of these, three proteins were cleaved upon expression of the viral protease to generate fragments of the expected size (PLSCR1, PLEKHA4, and WDR33; Fig. 5E and Supplementary Table S8). Of note, while WDR33 was predicted to harbor two potential cleavage sites, only a single cleavage event was observed. Treatment with a specific 3CD^pro^ inhibitor, rupintrivir ^31^, blocked the cleavage of these proteins, indicating the effect was due to the viral protease (Fig. 5D). In contrast, five of the proteins were found to not be cleaved upon 3CD^pro^ expression, suggesting additional determinants are involved in the cleavage of host factors (Fig. S6). Hence, our approach correctly identified 30% of the predicted cleavage sites (3 of the 9 different cleavage sites), indicating a strong enrichment of cellular targets of the 3CD^pro^ in the dataset.

## CONCLUSIONS

The picornavirus capsid is a highly complex structure that plays key roles in viral biology and pathogenesis. In the current study, we employ a comprehensive approach to define the effects of single amino acid mutations in the CVB3 capsid, measuring the effects of >90% of all possible mutations. We find that most mutations in the capsid are deleterious, with very few mutations showing higher fitness than the WT sequence (1.2% of all mutations). Similar results have been reported in other non-enveloped capsid proteins ^32–34^ as well as non-capsid viral proteins ^17,35–41^. In light of these results, it is likely that the large population sizes of RNA viruses help maintain viral fitness in the face of high mutation rates and strong mutational fitness costs.

Investigation of the factors that influence MFE in the capsid revealed a strong correlation with various structural and functional attributes. These included computationally predicted effects on stability and aggregation propensity, secondary structure, and surface exposure (Fig. 2). Surprisingly, we find that MFE can be predicted with relatively high accuracy using only five parameters: natural sequence variation, the identity of the original and mutant amino acid, the predicted effect on protein stability, and relative solvent accessibility (Fig. 3). A recent study examined the ability of 46 different variant effect prediction tools to predict MFE from 31 different DMS datasets of both viral and non-viral proteins ^42^. Overall, viral proteins showed the lowest predictability (Spearman’s correlation of <0.5). In contrast, we were able to predict MFE using a random forest model using these above-mentioned five parameters with an accuracy similar to the best prediction obtained in this analysis for any viral or non-viral protein (Pearson’s r= 0.73; Spearman’s ρ = 0.73; Fig. 3B). Interestingly, SNAP2 ^43^, a neural network-based classifier of mutational effects that was shown to correlate well with MFE in other studies ^42,44,45^, correlated poorly with our data (R^2^ = −0.26). Overall, considering the relative conservation of capsid structure in picornaviruses as well as the availability of both capsid sequences and high-resolution structures for numerous members of this family, it is likely that these findings can be extrapolated to additional picornaviruses.

Incorporating site-specific amino acid preferences obtained from our DMS results into phylogenetic models was found to significantly improve model accuracy. This has been observed in DMS studies with other RNA viruses ^24,39,46^, and indicate that our laboratory-measured MFE capture additional information that cannot be obtained from sequence analysis alone. In addition, this approach allowed us to assess which sites show differential selection patterns as a result of the distinct environments encountered in nature and the laboratory. As expected, pressure from the adaptive immune system was found to be the major difference between these environments, with residues in antibody neutralization sites showing higher differential selection compared to other sites in the capsid (Fig. 4A). Moreover, the sites showing the highest degree of differential selection were found in known antibody neutralization sites (Fig. 4B-D). However, why these particular residues within antibody neutralization sites show differential selection, while others do not, remain to be elucidated. It has been shown that one, or a few, sites within antibody binding regions can have strong effects on escape from antibody neutralization ^47^, potentially explaining these findings. Interestingly, while the top three sites showing differential selection were in antibody neutralization sites, the mutation showing the fourth-highest differential selection was found in the HI loop of VP1. While not classically considered an antibody epitope, this loop has been shown to interact with an antibody fragment in the picornavirus coxsackievirus A6 ^48^, is known to mediate receptor binding in different picornaviruses ^49,50^, and to interact with host cyclophilin A to facilitate uncoating ^15^. Whether these factors or others are responsible for the observed differential selection remains to be elucidated.

The CVB3 capsid encodes the information for directing myristoylation, protease cleavage, and interaction with host factors. We took advantage of our data to examine the sequence specificity and mutational tolerance of several known capsid encoded motifs. First, we examined the amino acid preferences of the CVB3 capsid myristoylation motif. We observe a strong correlation with the canonical myristoylation pattern (Prosite pattern PDOC00008), although with greater intolerance to mutations in three of the six residues in the capsid (Fig. S5). This is likely to stem from additional constraints imposed by capsid structure. On the other hand, we examined the amino acid preference of a conserved motif in VP1 that is required for 3CD^pro^-mediated cleavage of picornavirus capsids ^26^. Our data showed a higher cost to mutation in this motif relative to other capsid positions (Fig. S5), highlighting its importance for capsid function. Finally, we examined the sequence preferences surrounding the two 3CD^pro^ cleavage sites. We find a strong dependence on the cleavage site residues (positions P1 and P1’; Fig. 5) and to a lesser degree position P4, with large variation in the sequence preferences across the remaining positions between the two cleavage sites. Overall, our experimentally measured MFE are congruent with existing information regarding the sequence preferences of the examined capsid motifs, yet provide in-depth insights into sequence specificity that cannot be obtained from examining natural sequence variation.

Finally, we used the amino acid preferences observed in 3CD^pro^ cleavage sites within the capsid to query the human genome for potential cellular targets of this protease (Fig. 5D). Using this approach, we identify 746 cytoplasmic proteins that harbor a potential 3CD^pro^ target sequence, including 11 proteins previously shown to be cleaved by different picornavirus 3C proteases. We then validated our approach using eight proteins, comprising nine predicted cleavage sites. Six of the predicted cleavage sites were not affected by 3CD^pro^ expression (Fig. S6). On the other hand, three proteins were observed to be specifically cleaved by the viral protease (Fig. 5E): WD Repeat Domain 33 (WDR33), an important factor for polyadenylation of cellular pre-mRNAs ^51^ that has been shown to act as a restriction factor during influenza infection ^52^; the interferon-induced protein Phospholipid scramblase 1 (PLSCR1), which is involved in the replication of numerous viruses, likely due to its ability to enhance the expression of certain interferon-stimulated genes ^53^; and the interferon-induced Pleckstrin Homology Domain Containing A4 (PLEKHA4), a plasma membrane-localized signaling modulator ^54^ that is currently not known to play a role in viral infection. Overall, our approach correctly predicts 30% of the identified cleavage sites. It is likely that incorporating additional selection criteria, such as accessibility of the cleavage peptide in the folded structure, can be used to further reduce false positives. Nevertheless, extrapolating our validation results to the larger dataset suggests >200 new host targets of the protease are identified, many of which could play key roles in viral biology and pathogenesis.

## ACKNOWLEDGMENTS

The authors would like to thank Dr. Javier O. Cifuentes for help with the interpretation of antibody neutralization sites and Drs. Santiago Elena and Tzachi Hagai for critical reading of the manuscript. In addition, the authors would like to acknowledge the use of the Principe Felipe Research Center (CIPF) server which was co-financed by the European Union through the Operativa Program of the European Regional Development Fund (ERDF/FEDER) of the Comunitat Valenciana 2014-2020.

## AUTHOR CONTRIBUTIONS

FM, VL, and RG designed and performed the experiments. FM, VL, OT, AS and RG analyzed the data. FM, VL, AS, and RG wrote the manuscript. RG acquired funding.

## METHODS

### Viruses, cells, and plaque assays

HeLa-H1 (CRL-1958) and HEK293 (CRL-1573) cells were obtained from ATCC. All work with CVB3 was based on the Nancy infectious clone (kind gift of Dr. Marco Vignuzzi, Institute Pasteur). Cells were cultured in culture media (DMEM with 10% heat-inactivated FBS, Pen-Strep, and L-Glutamine) with FBS concentrations of 2% during infection. For plaque assays, serial dilutions of the virus were used to infect confluent HeLa-H1 cells in 6 well plates for 45 minutes, followed by overlaying the cells with a 1:1 mixture of 56°C 1.6% Agar (Arcos Organics 443570010) and 37°C 2x DMEM with 4% FBS. Two days later, plates were fixed with formaldehyde (2% final concentration) after which the agar was removed and the cells stained with crystal violet to visualize plaques.

### Deep mutational scanning (DMS)

The infectious clone was modified by site-directed mutagenesis to remove an XhoI site present in the capsid region (P1) and introduce an XhoI site at position 692 as well as a Kpn2I site at position 3314, generating a pCVB3-XhoI-P1-Kpn2I clone ^55^. In addition, a pCVB3-XhoI-ΔP1-Kpn2I plasmid was generated by replacing the region between the XhoI and Kpn2I sites in pCVB3-XhoI-P1-Kpn2I with a short linker. To generate the template for DMS, the capsid region was amplified by PCR from pCVB3-XhoI-P1-Kpn2I with Phusion polymerase (Thermo Scientific) and primers HiFi-F (CTTTGTTGGGTTTATACCACTTAGCTCGAGAGAGG) and HiFi-R (CCTGTAGTTCCCCACATACACTGCTCCG) and gel purified (Zymoclean Gel DNA Recovery Kit). Primers spanning the full coding region of the capsid region were designed using the CodonTilingPrimers software from the Bloom lab (https://github.com/jbloomlab/CodonTilingPrimers) with the default parameters and synthesized by IDT (Supplementary Table S1). These primers were used to perform the mutagenesis PCR on the capsid template together with the HiFi-F or HiFi-R primers in triplicate following published protocols ^56^ with the exception that 10 rounds of mutagenesis were performed for libraries 1 and 2, while a second round of 7 mutagenesis cycles was performed for library 3 to increase the number of mutation per clone. The products were gel purified and ligated to an XhoI and Kpn2I digested and gel purified pCVB3-XhoI-ΔP1-Kpn2I using NEBuilder^®^ HiFi DNA Assembly reaction (NEB) for 25 minutes. Mutagenesis efficiency was evaluated by the transformation of the assembled plasmids into NZY5α competent cells (NZY Tech), Sanger sequencing of 18-23 clones per library, and mutation analysis using the Sanger Mutant Library Analysis script (https://github.com/jbloomlab/SangerMutantLibraryAnalysis). Subsequently, the assembled plasmid reactions were purified using a Zymo DNA Clean & Concentrator-5 kit (Zymo Research) and used to electroporate MegaX DH10B T1R Electrocomp cells (ThermoFisher) using a Gene Pulser XCell electroporator (BioRad) according to the manufacturer’s protocol. Cells were then grown overnight in a 50 mL liquid culture at 33°C and DNA purified using the PureLink HiPure plasmid midiprep kit (Invitrogen). Transformation efficiency was estimated by plating serial dilutions of the transformation on agar plates. In total, 4.44×10^5^, 1.46×10^5^, and 2.19×10^5^ transformants were obtained for lines 1, 2, and 3, respectively. Viral genomic RNA was then transcribed from SalI linearized, gel-purified full-length plasmids using the TranscriptAid T7 kit (ThermoScientific), and four electroporations were performed using 4×10^6^ HeLa-H1 cells in a 4mm cuvette in 400μL of calcium and magnesium-free PBS using with 8μg of RNA in a Gene Pulser XCell (BioRad) set to 240V and 950uF. Electroporated cells were then pooled, and one fourth was cultured for 9 hours to produce the passage 0 virus (P0). Following three freeze-thaw cycles, 2×10^6^ plaque-forming units (PFU) were used to infect a 90% confluent 15cm plate in 2.5mL of infection media for 1 hour. Cells were then washed with PBS and incubated in 12 mL of infection media for 9 hours. Finally, cells were subjected to 3 freeze-thaw cycles, debris removed by centrifugation at 500xg and the supernatants collected to generate P1 virus stocks. All infection produced > 2.38×10^6^ PFU in P0 and > 1.2×10^7^ PFU in P1 as judged by plaque assay.

### NGS analysis

Libraries were prepared following published protocols ^57^ and each library was run on a Novaseq6000 2×150 at a maximum of 30G per lane to reduce potential index hopping. Reads trimming was performed using fastp ^58^ (command: -max_len1 150 --max_len2 150 --length_required 150 −x −Q −A), unsorted bam files were generated from fastq files using Picard tools FastqToSam (version 2.2.4) and merged into a single bam using the cat command of Samtools (version 1.5). The duplex pipeline was then implemented (https://github.com/KennedyLabUW/Duplex-Sequencing/UnifiedConsensusMaker.py) using the UnifiedConsensusMaker.py script and a minimum family size of 3, a cutoff of 0.9 for consensus calling, and an N cutoff of 0.3. The single-stranded consensus files (SSCS) were then aligned using BWA mem (version 0.7.16), sorted using Samtools, size selected to be 133 bp long using VariantBam ^59^, unaligned reads were discarded (Samtools view command with −F 4), and the resulting bam file indexed with Samtools. Subsequently, fgbio (http://fulcrumgenomics.github.io/fgbio/; version 1.1.0) was used to hard-clip 10 bp from each end and upgrade all clipping to hard-clip (−c Hard --upgrade-clipping true --read-one-five-prime 10 --read-one-three-prime 10 --read-two-five-prime 10 --read-two-three-prime 10). Variant bam was then used to keep all reads that were between 50-150bp, well-mapped, and had either no indels and less than 5 mutations (command –r {“”:{“rules”:[{“ins”:[0,0],“del”:[0,0],“nm”:[0,4], “mate_mapped”:true,“fr”:true,“length”:[50,150]}]}}’). Finally, the codons in each read were identified using the VirVarSeq ^60^ Codon_table.pl script using a minimum read quality of 20. A custom R script was then used to generate a codon counts table for each codon position by eliminating all codons containing ambiguous nucleotides and codons with a strong strand bias (StrandOddsRatio > 4), as well as all codons that are reached via a single mutation (available at https://github.com/RGellerLab/CVB3_Capsid_DMS). Amino acid preferences and mutational fitness effects were determined using DMStools2 ^19^ with the Bayesian option and the default settings.

### Structural analyses

The crystal structure PDB:4GB3 ^61^ was used for all structural analysis. The effects of mutations on aggregation were determined using TANGO version 2.3.1 ^22^ using the default settings and the effect on stability on the monomer and pentamer was determined using FoldX 4 ^21^ using the default settings. For the latter, the pentamer subunits were renamed to unique letters, all mutations between the reference sequence and the structure sequence were introduced using the BuildModel command, the structure was optimized using the RepairPDB command 5 or 10 times for the pentamer or monomer, respectively, and then the effects of the mutations were predicted using the BuildModel command (modified PDB files can be found at https://github.com/RGellerLab/CVB3_Capsid_DMS). Secondary structure and RSA were obtained from DSSP (http://swift.cmbi.ru.nl/gv/dssp/) using the dms_tools2.dssp function of dms_tools2, while interface, surface, and core residues as well as residue contact number, and presence in the two, three, and five-fold axes were obtained from ViprDB (http://viperdb.scripps.edu/) ^62^. Distance from the center was calculated with Pymol using the Distancetoatom.py script on the monomer or pentamer.

### Generation and evaluation of CVB3 capsid mutants

The PCR of the capsid region used as a template for DMS was phosphorylated and cloned into a SmaI digested pUC19 vector for use in the mutagenesis reactions (pUC19-HiFi-P1). For each mutant, non-overlapping primers containing the mutation in the middle of the forward primer were used to introduce the mutation with Phusion polymerase, followed by DpnI (Thermo Scientific) treatment, phosphorylation, ligation, and transformation of chemically competent bacteria. Successful mutagenesis was verified by Sanger sequencing. Subsequently, the capsid region was subcloned into pCVB3-XhoI-ΔP1-Kpn2I using XhoI and Kpn2I sites. Plasmids were then linearized with MluI and 2μg of plasmid was transfected into 5×10^5^ HEK293 cells together with a plasmid encoding the T7 polymerase ^63^ (Addgene 65974) using calcium phosphate. Briefly, an equal volume of 2x HBS (274mM NaCl, 10mM KCl, 1.4mM Na_2_HPO_4_) was added dropwise to DNA containing 0.25M CaCl_2_ while mixing, incubated 15 minutes at RT, and then added dropwise to cells. Following 48 hours, passage 0 (P0) virus was collected and titered by plaque assay. From this, 10^5^ PFU were used to infect 90% confluent 6 well HeLa-H1 cells (MOI 0.1) for 1 hour at 37°C, after which the cells were washed twice with PBS and 2mL of infection media added. Cells were then incubated until CPE was observed. Emerging viral populations were titered by plaque assay and the capsid region sequenced to ensure no compensatory mutations or reversions arose during replication. The fitness of these mutants was then tested by direct competition with a marked reference virus using a Taqman RT-PCR method ^20^. Briefly, in quadruplicates, confluent HeLa-H1 cells in a 24 well plate were infected with 200μL of a 1:1 mixture of 4×10^3^ PFU (MOI 0.01) of the test and marked reference viruses for 45 minutes. Subsequently, the inoculum was removed, the cells were washed twice with PBS, 200 μL of infection media was added, and the cells were incubated for 24 hours at 37°C. Finally, cells were subjected to 3 freeze-thaw cycles, debris removed by centrifugation at 500xg, the supernatants collected and treated with 2uL of RNase-Free DNaseI (ThermoFisher) for 15 minutes at 37°C, and viral RNA extracted using the *Quick*-RNA™ Viral Kit (Zymo Research), eluting in 20μl. Quantification of the replication of each mutant versus the reference was performed using Luna^®^ Universal Probe One-Step RT-qPCR kit (New England BioLabs) containing 3uL of total RNA, 0.4μM of each qPCR primers and 0.2μM of each probe. The standard curve was performed using 10-fold dilutions of RNA extracted from 10^7^ PFU of wild-type and reference viruses. All samples were performed with three technical replicates. The relative fitness (W) of each mutant versus the common marked reference virus was calculated using the formula W = [R(t)/R(0)]^1/t^, where R(0) and R(t) represents the ratio of the mutant to the reference virus genomes in the initial mixture used for the infection and after 1 day (t=1), respectively ^20,64^.

### Sequence variability and phylogenetic analyses

Amino acid variability was assessed using Shannon entropy. Briefly, all available, non-identical, full-genome CVB3, CVB, or Enterovirus B sequences were downloaded from Virus Pathogen Resource ^65^ (www.viprbrc.org) and codon-aligned using the DECIPHER package in R (available at https://github.com/RGellerLab/CVB3_Capsid_DMS). All alignment positions not present in our reference strain were removed, and a custom R script was used to calculate Shannon entropy. For phylogenetic and differential selection analyses, PhyDMS was run using the default settings on an alignment of CVB3 genomes that was processed with the phydms_prealignment module and using the average preferences from the three DMS replicates.

### Identification of 3CD^pro^ cleavage sites in the human proteome

The amino acid preferences (the relative enrichment of each amino acid at each position standardized to 1) was used to generate in silico 1000 peptides spanning the 10 amino acid region surrounding each cleavage site using a custom R script (available at https://github.com/RGellerLab/CVB3_Capsid_DMS). Specifically, for each peptide position, 100 peptides were generated that encoded each amino acid at a frequency corresponding to its preference observed in the DMS results, with the remaining positions unchanged. The resulting 1000 peptides from each cleavage site were uploaded to PSSMSearch ^66^ (http://slim.icr.ac.uk/pssmsearch/) using the default setting (psi_blast IC). Results were filtered to remove proteins indicated to be secreted, lumenal, or extracellular in the Warnings column. To test whether proteins were cleaved by the viral 3CD protease, the corresponding region was PCR amplified from the Nancy infectious clone (primers 3C-For: TATTCTCGAGACCATGGGCCCTGCCTTTGAGTTCG and 3D-Rev: TATTGCGGCCGCCTAGAAGGAGTCCAACCATTTCCT) and cloned into the pIRES plasmid (Clonetech) using the restriction sites XhoI and NotI (pIRES-3CD^pro^). For analysis of fusion proteins, HEK293 cells were transfected with GFP-PLEKHA4 (kind gift of Dr. Jeremy Baskin, Cornell University), GFP-PLSCR1 (kind gift of Dr. Serge Benichou, Institut Cochin), FLAG-NLCR5 (Addgene #37521), HA-ZC3HAV1 (Addgene #45907), or the control plasmid FLuc-eGFP (Addgene #90170) together with the pIRES-3CD^pro^ plasmid using Lipofectamine 2000. Following 24 hours, proteins were collected by lysing in lysis buffer (50mM TRIS-HCl, 150mM NaCl, 1% NP40 and protease inhibitor cocktail [Complete Mini EDTA-free, Roche]) and subjected to western blotting with the corresponding antibody (anti-GFP, Santa Cruz sc-9996; Anti FLAG, Santa Cruz sc-166335; anti-HA, Santa Cruz, sc-7392). For analysis of endogenous proteins, 3CD^pro^ was expressed for 48 hours before cell lysis, and western blotting using antibodies against WDR33 (Santa Cruz sc-374466), TSG101 (Santa Cruz sc-136111), GAK (Santa Cruz sc-137053), and MAGED1 (Santa Cruz sc-393291). When indicated, the 3C^pro^ inhibitor rupintrivir (Tocris Biosciences) was added at a concentration of 2 μM for the last 24 hours before collection. The predicted molecular weight of cleaved fragments was calculated using the mw function of the Peptides R package (version 2.4.2).

### Statistical analyses

All statistical analyses were performed in R and were two-tailed. For random forest prediction, the R RandomForest package (version 4.6-14) was employed using the default setting with an mtry of 10, and for the linear model, the formula lm(MFE ~ enterovirus B entropy + WT amino acid * mutant amino acid + predicted effect of mutations on stability in the pentamer + relative surface exposure) was used (available at https://github.com/RGellerLab/CVB3_Capsid_DMS). Sequence logoplots were producing using Logolas ^67^.

### Data availability

Unaligned bam files have been uploaded to SRA (Accession SAMN15437545-SAMN15437555; SRA 15437545-15437555). The scripts and data required to obtain the codon count tables for all samples, to perform the random forest and linear model predictions, to generate the peptides for use with PSSMsearch, as well as the sequence alignments and modified structure files for FoldX analysis can be found on Github (https://github.com/RGellerLab/CVB3_Capsid_DMS).

---

**Supplementary Figure S1.**
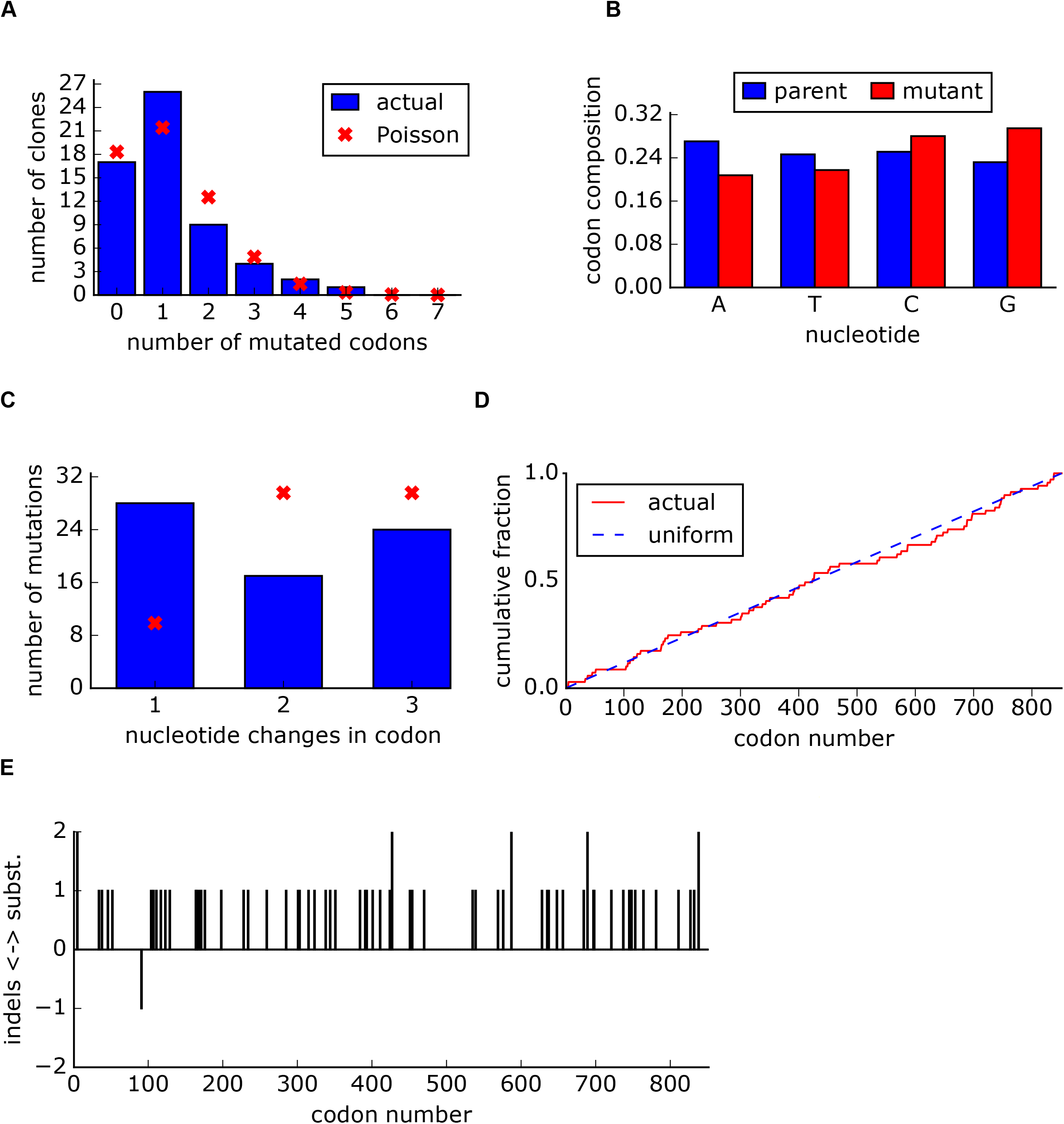
Sanger analysis of DMS libraries. **A.** The number of mutated codons per clone. **B.** Original and mutated base for each mutation. **C.** The number of nucleotide changes per codon. **D.** Cumulative fraction of mutations versus the codon position. **E.** Location of both mutations and indels across the capsid sequence.

**Supplementary Figure S2.**
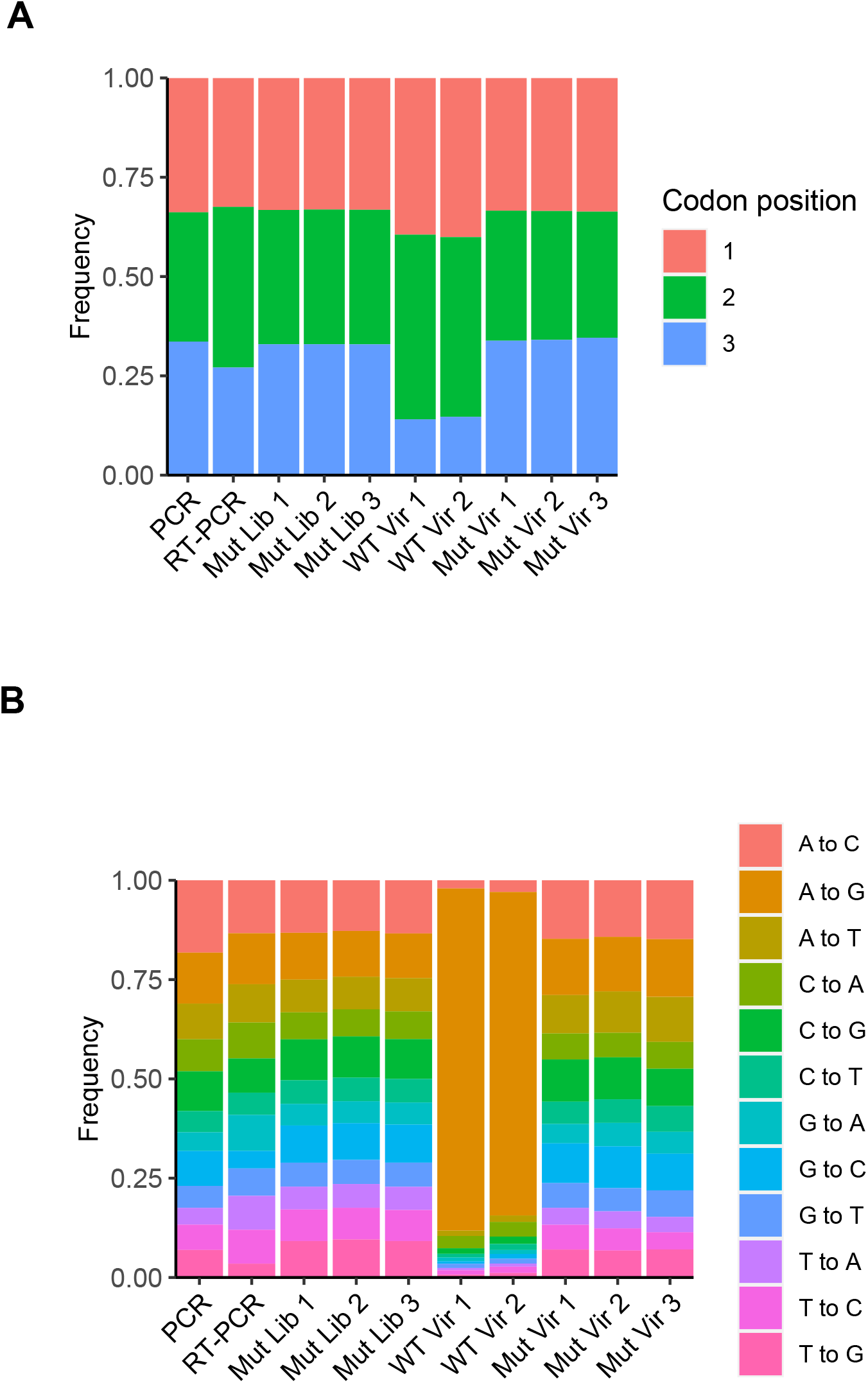
Results of high-fidelity duplex sequencing. **A.** The relative frequency of the mutated base within each mutated codon. **B.** The relative frequency of each mutation type.

**Supplementary Figure S3. Correlation of amino acid preferences observed in experimental replicates.** Hexagonal bin plots showing the correlation of amino acid preferences between the three experimental replicates. Spearman’s correlation coefficient and p-value are shown above each plot.

**Supplementary Figure S4.**
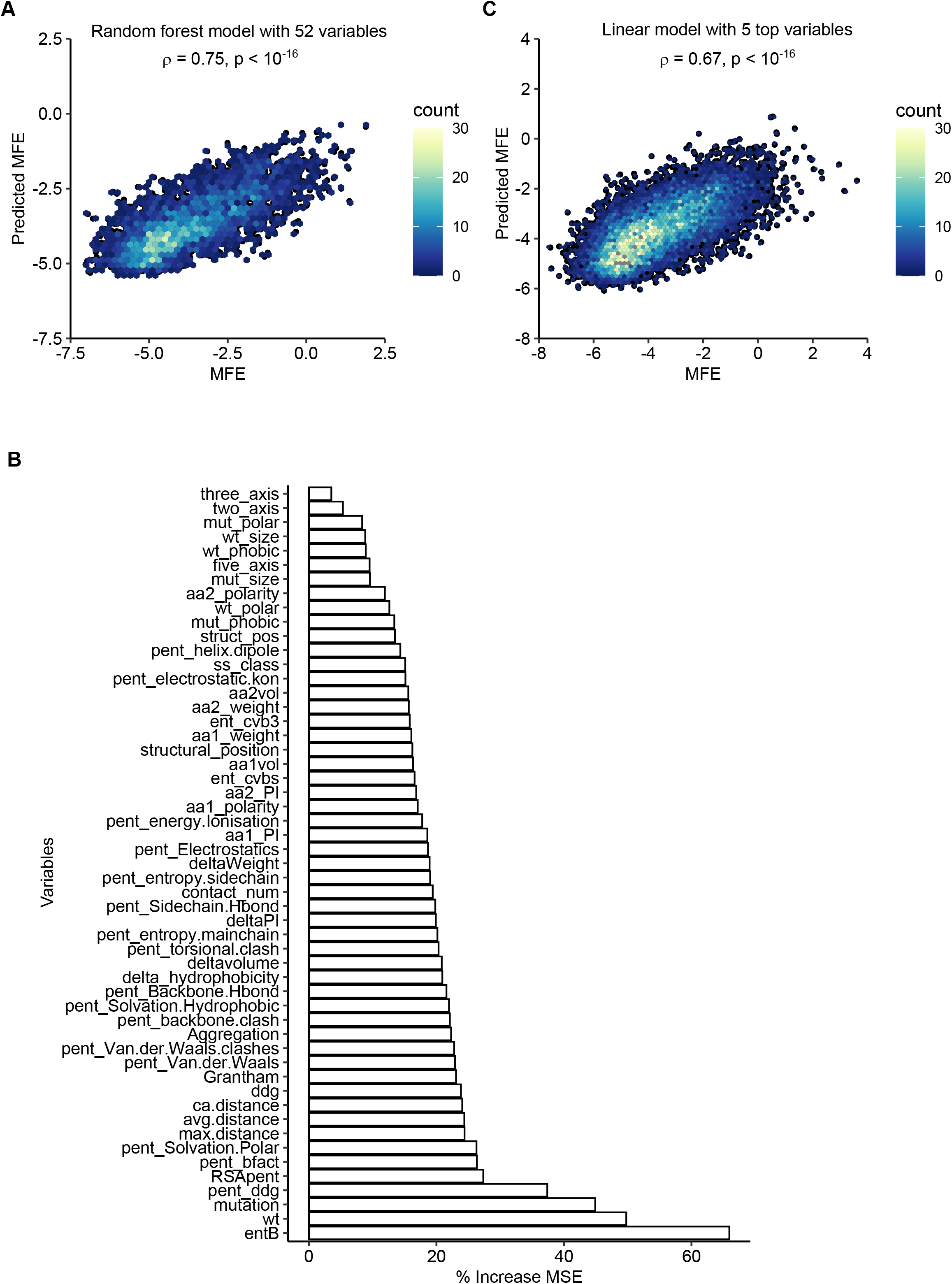
Prediction of mutational fitness effects using random forest or linear models. **A.** Hexagonal bin plot showing the correlation between actual and predicted MFE derived from a random forest model using all 52 variables. The model was trained on 70% of the data and tested on the remaining 30% of the data (shown). **B.** Variable importance obtained from the random forest model. **C.** Linear model using the top five parameters of the random forest model. See supplementary Table S6 for parameter description.

**Supplementary Figure S5.**
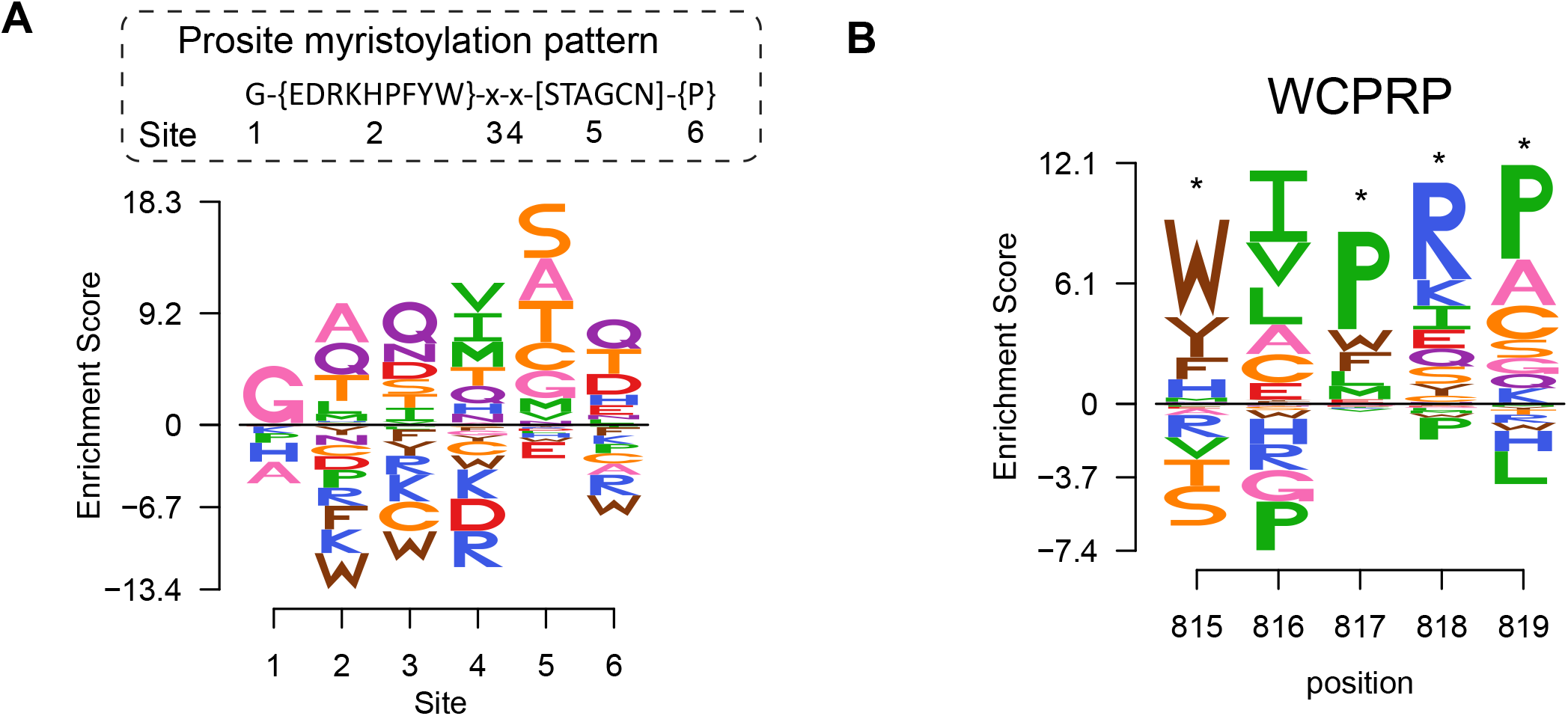
Sequence preferences of capsid encoded motifs. **A.** Amino acid preferences of the CVB3 myristoylation motif. The canonical Prosite myristoylation motif is indicated above, with curly brackets indicating disfavored amino acids and square brackets indicating tolerated amino acids. **B.** WCPRP motif required for 3CD^pro^ cleavage of P1. Asterisks indicate analogous positions in FMDV shown to be essential for viability ^26^.

**Supplementary Figure S6. Evaluation of select hits identified as potential 3CD^pro^ target proteins.**

Western blots of cells transfected with 3CD^pro^ and probed for the indicated endogenous protein, or cotransfected with 3CD^pro^ and the indicated fusion protein and blotted for the tag. Each experiment was performed twice. When indicated, the 3C^pro^ inhibitor rupintrivir was added.

